# Myeloid-derived Suppressor Cells Mitigate Inflammation in Periodontal Disease

**DOI:** 10.1101/2024.12.21.629927

**Authors:** Raza Ali Naqvi, Araceli Valverde, Salvador Nares, Thomas E. Van Dyke, Afsar R. Naqvi

## Abstract

Myeloid derived suppressor cells (MDSCs) are a heterogeneous population of immature, immune suppressive myeloid cells. However, their role in periodontal disease (PD), a microbially induced inflammatory oral disease, remains understudied. Here we show that gingiva (gums) from PD patients exhibit significantly higher levels of MDSC markers including Granulocytic (G)-MDSC and Monocytic (M)-MDSC subsets as well as CD4^+^ T cells and CD19^+^ B cells. Gingival MDSC subsets exhibit potent immunoregulatory activity as marked by attenuated autologous CD4+ T cells proliferation and IFNγ production. In a murine model of ligature-induced periodontitis (LIP), we noticed time-dependent gingival MDSC infiltration, which correlates with CD4^+^ T cell and CD19^+^ B cell infiltration. To test whether MDSC confer immunoregulatory function *in vivo*, we adoptively transferred G-MDSCs and M-MDSCs in mice. Interestingly, we observed significant reduction in inflammatory marker expression (IL6, TNF-α, and IL-1β), infiltration of CD4^+^ T cells, and concomitant increase in MDSC-derived immune suppressive molecules, ARG1 and IL-10, and CD4^+^CD25^+^FoxP3^+^ Tregs compared to mock. Conversely, depletion of MDSC using anti-Gr1 antibody resulted in marked induction of periodontal inflammation, reduced Treg population, and significantly higher alveolar bone loss. These findings, for the first time, suggest an anti-inflammatory and osteoprotective function of MDSCs in PD and offer a promising target to treat unresolved periodontal inflammation.

## INTRODUCTION

Periodontal diseases (PD) are a group of infectious, inflammatory disorders of the supporting structures of teeth, affecting about 750 million people worldwide.^1-4^ PD is initiated by a dysbiotic oral biofilm and is followed by an aberrant host immune response.^2-5^ Moreover, PD is associated with systemic diseases including neurological diseases, rheumatoid arthritis, obesity, cardiovascular diseases, diabetes, and others.^6-9^ Therefore, modulating periodontal inflammation has the potential to improve overall health. Current periodontal treatment involves mechanical plaque biofilm/calculus (tarter) debridement or surgical intervention but have yielded limited success.^3,4^ Development of regulatory endogenous molecules that can modulate periodontal inflammation is of high clinical significance.

An overt and persistent proinflammatory microenvironment is the primary determinant of PD progression.^10^ Multiple lines of evidence, including studies from our lab, clearly demonstrate the accumulation of inflammatory immune cells in periodontitis. Firstly, higher accumulation of myeloid macrophages (Mφ) / dendritic cells (DC), T cells and B cells occurs in diseased gingival biopsies.^5,11,12^ Secondly, copious levels of immunoglobulins, cytokines and inflammation-associated genes are detected in inflamed human gingiva.^13,14^ Thirdly, nonresolving inflammation may persist/relapse even after routine periodontal therapy.^15^ These observations clearly highlight perturbed immune responses. Therefore, modulation of the pro-inflammatory immune microenvironment via cell therapy of immune regulatory cells capable of maintaining immune homeostasis may serve as a promising therapeutic strategy to treat PD or mitigate disease progression.

Myeloid-derived suppressor cells (MDSCs) are a heterogeneous immune regulatory population of immature myeloid cells^16,17^ exerting immune suppression via increased production of arginase-1 (ARG1), inducible nitric oxide synthetase (iNOS) activity, and the induction of T-cell apoptosis. Based on their origin, MDSC are broadly classified into two subsets: monocytic (M-MDSC) and granulocytic (G-MDSC), both capable of exhibiting potential immunoinhibitory effects.^17-19^ MDSCs have been shown to have inhibitory effects in mouse tumor models and human cancers.^20^ Importantly, human MDSCs are characterized by the co-expression of CD11b^+^ and CD33^+^ and low levels of HLA-DR (Class II MHC). However, CD11b and Gr-1 are identified as markers for murine derived MDSCs in HLA-DR low population of cells.^16,21^ Recently, early state MDSCs (e-MDSCs) have been described as cells that co-express CD33 and CD11b in HLA-DR^lo^ cells, and lack the myeloid markers CD14 and CD66b.^22,23^ Various reports demonstrate that MDSCs can serve as an effective therapy for various autoimmune diseases including inflammatory bowel disease (IBD), experimental autoimmune rheumatoid arthritis, systemic lupus erythematosus (SLE),^24-26^ and maintenance of immune tolerance during organ transplantation.^27,28^ However, the functional role of gingival MDSCs in PD pathobiology remains less explored.

In this report, we characterized MDSC infiltration in subjects with PD and validated their infiltration kinetics in a murine model of ligature induced periodontitis (LIP). Data from both experiments confirm significant homing of MDSCs in gingiva; however, higher CD4^+^ T cells and CD19^+^ B cells in gingiva suggest an inability of MDSC to control periodontal inflammation. We hypothesize that a lack of adequate quantities of functional MDSCs could contribute to PD progression and might lead to unresolved inflammation. Using adoptive transfer and functional depletion experiments in the LIP model, we demonstrated that MDSCs perform immunoregulatory functions *in vivo*.

## RESULTS

### Gingival infiltration of M-MDSC and G-MDSC occurs in periodontal disease

In PD, excessive accumulation of pro-inflammatory myeloid and lymphoid cells suggests an impairment of regulatory immune cell function.^5,11,12,29^ The role of myeloid immunoregulatory cells in PD, however, is less explored. We asked whether MDSC, a key innate immunoregulatory population, are present in inflamed gingiva. In this pursuit, gingiva was collected from healthy and PD subjects to characterize MDSC by multiparametric flow cytometry. **Fig. 1A, B** describe the evaluation of two distinct populations: M-MDSC (CD11b^+^CD33^+^HLA-DR^lo^CD14^+^) and G-MDSC (CD11b^+^CD33^+^HLA-DR^lo^CD14^-^) in the gingiva of healthy and periodontitis patients. We observed significantly higher proportion of both CD11b^+^CD33^+^HLA-DR^lo^ cells (7.2 ± 1.2 % *vs* 2.2 ± 0.45%; P<0.005) in inflamed gingiva compared to healthy gingiva (**Fig. 1C**). Furthermore, RT-qPCR data showed a significant increase in the expression of MDSC associated markers including CD33 (fold change: 14.1 ± 1.34; P<0.001) (**Fig. 1D**), and pro-inflammatory cytokine genes: IL-6 (fold change: 23.1 ± 3.20; P<0.001) and TNF-α (fold change: 16.3 ±2.02; P<0.001) in gingival biopsies of PD subjects compared to healthy controls (**Fig. 1C-E**). Together, these results demonstrate MDSC infiltration in inflamed gingiva.

**Figure 1.**
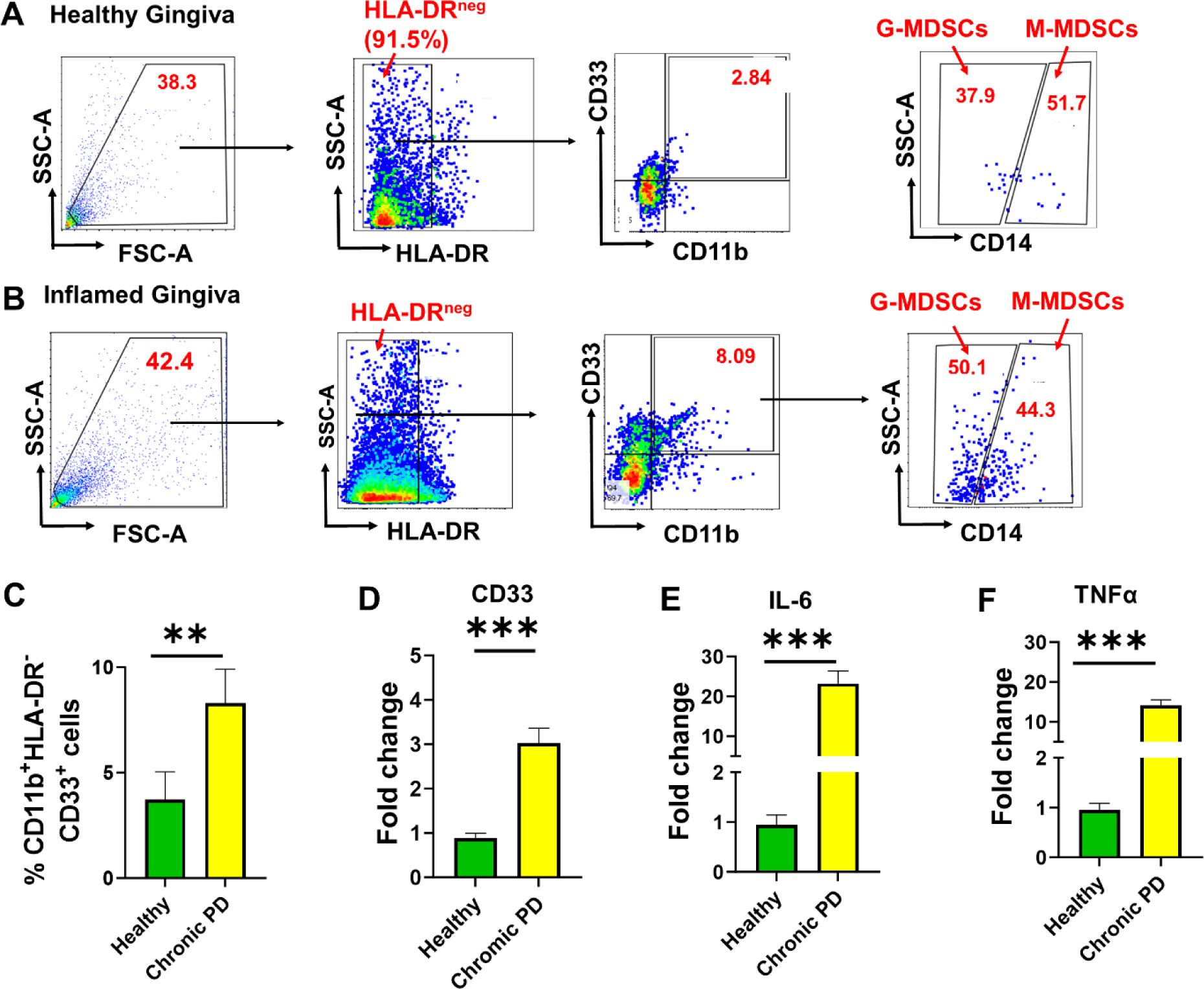
Higher numbers of MDSCs subsets infiltrate gingiva in periodontal disease. Gingiva collected from periodontally healthy and diseased human subjects (n=10/group) were examined for M-MDSCs and G-MDSCs subsets. Representative figure demonstrating M-MDSCs and G-MDSCs proportion in (A) healthy and (B) inflamed gingiva. (C) Comparison of total MDSC (HLA-DR^-^ CD11b^+^CD33^+^) population in gingiva of periodontally healthy and diseased subjects. RT-qPCR showing the expression of (D) CD33 (E) IL-6, and (F) TNF-α in human gingiva. Data is presented as mean ±S.D. Student’s t test was used to calculate the significance between healthy vs periodontitis gingiva samples. Student’s t-test was performed to evaluate the significance. *P< 0.05;**P< 0.01;***P< 0.001.

Chronic periodontal lesions are marked by infiltrating CD4^+^ T cells and CD19^+^ B cells.^32,33^ Consistent with this, we observed significantly higher proportions of both CD4^+^T cells (8.95 ± 1.13% *vs* 2.5 ± 0.87%; P<0.0005), which exhibit an inflammatory phenotype marked by IFNγ expression (CD4^+^IFNγ^+^ T cells; 7.1 ±1.6% *vs* 1.5 ± 0.82; P<0.005) (**Fig. 2 A,B)**. Moreover, we noticed a significantly higher percentage of CD19^+^ B cells (8.56 ± 1.86 % *vs* 1.8 ± 0.60; P<0.0005) in periodontally diseased verses healthy gingiva. Interestingly, we also observed significantly higher expression of IFNγ^+^ in B cells in gingiva from PD patients vs. healthy controls (2.4 ± 0.57 *vs* 1.2 ± 0.35%; P<0.005), which has not been previously reported (**Fig. 2C,D**).

**Figure 2.**
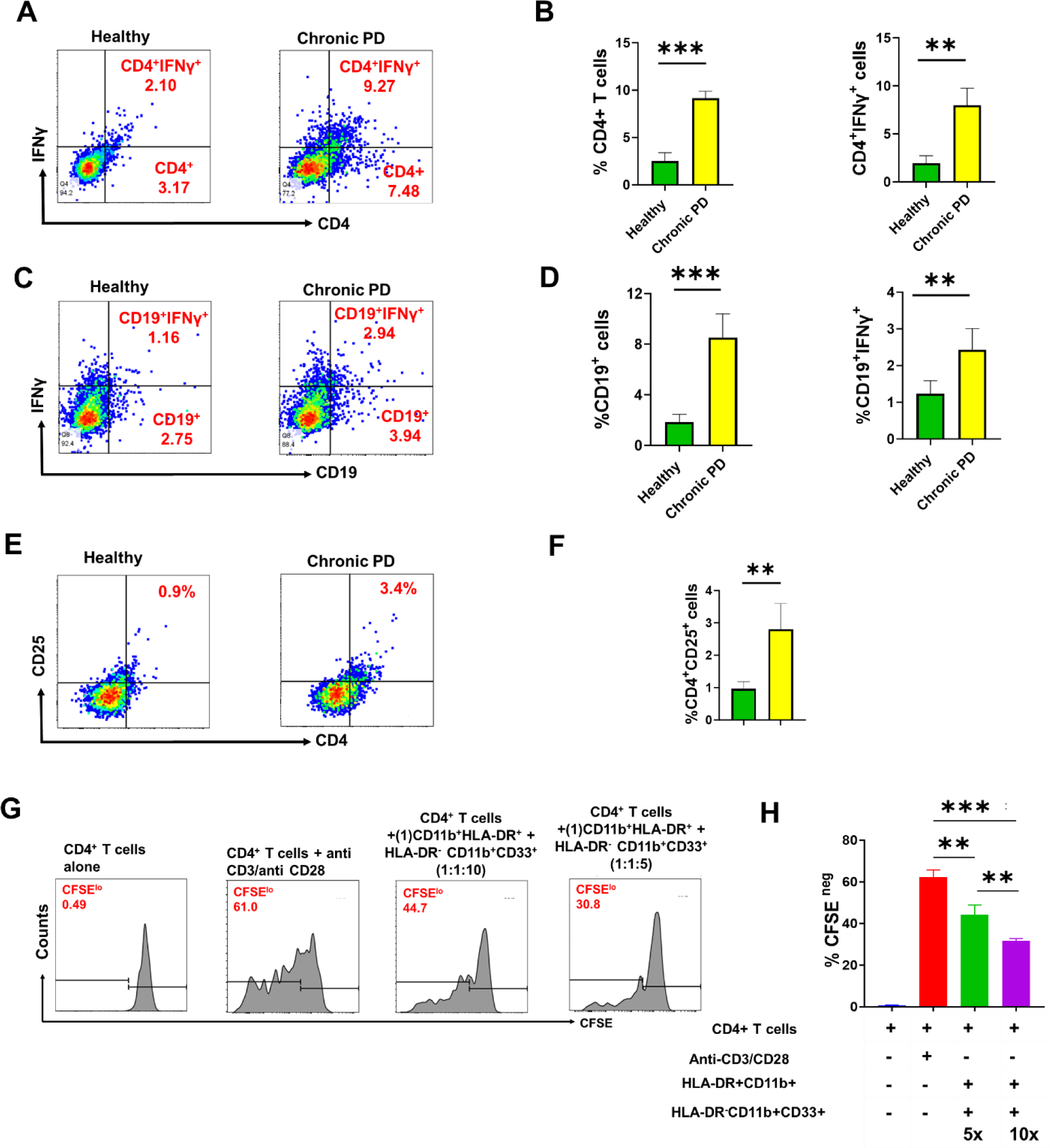
MDSCs infiltrated in inflamed gingiva exhibit immunoregulatory phenotype. Inflammatory T and B cells infiltrate periodontally diseased gingiva. (A) Scatter plot and (B) quantification of CD4^+^ T cells, and CD4^+^IFNγ^+^ T cells in periodontally healthy and inflamed human gingiva. (C) Scatter plots and (D) quantification of CD19+ B cells and CD19^+^IFNγ^+^ B cells in the gingiva of healthy control and periodontitis patients. (E) Scatter plots showing CD4^+^CD25^+^ T regulatory (Tregs) cells in healthy and PD subjects. (F) Quantitative analysis of CD4^+^CD25^+^ Tregs in healthy and PD shows higher numbers in inflamed gingiva. (G) MDSC isolated from inflamed gingiva suppress autologous T cell proliferation ex vivo. Sorted gingival MDSCs (CD11b^+^HLA-DR^-^CD33^+^), CD11b^+^HLA-DR^+^ (as antigen presenting cells) and CFSE labeled CD4^+^ T cells (n=5/group) were cultured in presence of 1mg/mL ovalbumin (as antigen) for 5 days. The numbers of MDSC and CD11b^+^HLA-DR^+^ were kept constant at 2x 10^5^, and CD4^+^ T cells were used in 1:5 and 1:10 ratios. At the end of the assay, cell proliferation was assessed by flow cytometric analysis. Histograms showing CD4 cell proliferation in different cocultures. (H) Bar graphs show percentages of proliferating CD4+ T cell population. Each bar in (B), (D), (F) and (H): represent Mean ±S.D. One-way ANOVA was used to calculate the significance in (H), while Student’s t-test was performed to evaluate the significance in (B), (D) and (F). *P< 0.05;**P< 0.01;***P< 0.001.

MDSCs enhance Treg quantities from a preexisting population of Tregs created by their secretory molecules ARG1, NO, ROS, IL-10, and TGF-β.^30,31^ Similar to MDSCs, we also observed a significant increase in CD4^+^CD25^+^FoxP3^+^ Tregs in gingiva of PD subjects compared to healthy controls (0.96% ±0.21 % vs 2.8 ± 0.79%; P<0.005) (**Fig. 2E,F**). These results indicate active lymphoid (Treg) and myeloid (MDSC) immunoregulatory pathways in PD.

Next, we asked whether infiltrated gingival MDSC elicit immunoregulatory functions. Sorted CD11b^+^CD33^+^ cells were co-cultured with CFSE labeled autologous CD4^+^ T cells obtained from inflamed gingiva of periodontitis subjects in various ratios. CFSE^lo^ cells were considered the proliferating population. We observed the following proportion of proliferating CD4+ T cells in various conditions: CD4^+^ T cells alone (0.56 ± 0.24 %), CD4^+^ T cells + anti CD3/CD28 beads (62.7 ± 3.59 %), CD4^+^ T cells + CD11b^+^HLA-DR^+^ + HLA-DR^-^CD11b^+^CD33^+^ (in 1:1:10) (44.16 ± 4.72) and CD4^+^ T cells + CD11b^+^HLA-DR^+^ + HLA-DR^-^CD11b^+^CD33^+^ (in 1:1:5) (31.66 ± 1.09). Compared to CD4^+^ T cells with anti CD3/CD28 beads, the proliferation of CD4^+^ T cells was significantly reduced in the presence of HLA-DR^-^CD11b^+^CD33^+^ in both 1:10 and 1:5 co-cultures. However, the proliferative inhibitory effect of MDSCs was more pronounced in 1:5 ratio co-cultures (**Fig. 2 G,H**).

Overall, these findings indicate marked infiltration as well as immune regulatory activities MDSCs in the gingiva of PD patients. However, MDSC infiltration fails to contain periodontal inflammation as reflected by higher inflammatory CD4^+^ T cell and CD19^+^ B cell counts.

### Infiltration of murine MDSCs during disease progression in a LIP model

To evaluate the homing of MDSCs during onset and progression of PD in a more controlled way, we examined their infiltration kinetics in a murine model of ligature induced periodontitis (LIP). This model was previously published by our group to dissect mechanisms underlying PD pathogenesis.^34,35^ Silk ligature placed between first and second maxillary molars induces microbial accumulation leading to induction of inflammation and subsequent alveolar bone loss by day 8. We evaluated the infiltration kinetics of MDSC in gingiva on day 2-, 4-, and 8-days post-ligature (DPL) placement. Previously, MDSCs in mice were described as myeloid cells containing CD11b and Gr-1 markers. Later and similar to humans, mice MDSCs were also categorized as PMN-MDSCs (expressing CD11b^+^Ly6G^+^) and M-MDSCs (CD11b^+^Ly6C^+^).^16,36^

Single-cell suspensions from murine LIP gingiva show remarkable infiltration of both MDSC subsets. We observed a time-dependent progressive increase in the infiltration of both MDSC subsets in murine gingiva. **Fig. 3A** depicts the schematics of the experiment. M-MDSC population increased from 1.5 ± 0.42% (4 DPL) to 3.7 ± 0.89% (P<0.001) at 8 DPL, while G-MDSC population spiked from 1.1 ± 0.43% (4 DPL value) to 5.2±1.2% by day 8 (P<0.001) (**Fig. 3B-E**). Furthermore, we also observed a significant increase in the gingival infiltration of CD4^+^T cells in the LIP murine model from 4DPL (6.8 ± 1.15%; P< 0.001) to 8 DPL (16.7 ± 3.0%; P<0.001) (**Fig. 3F,G**) indicating establishment of PD in this model.

**Figure 3.**
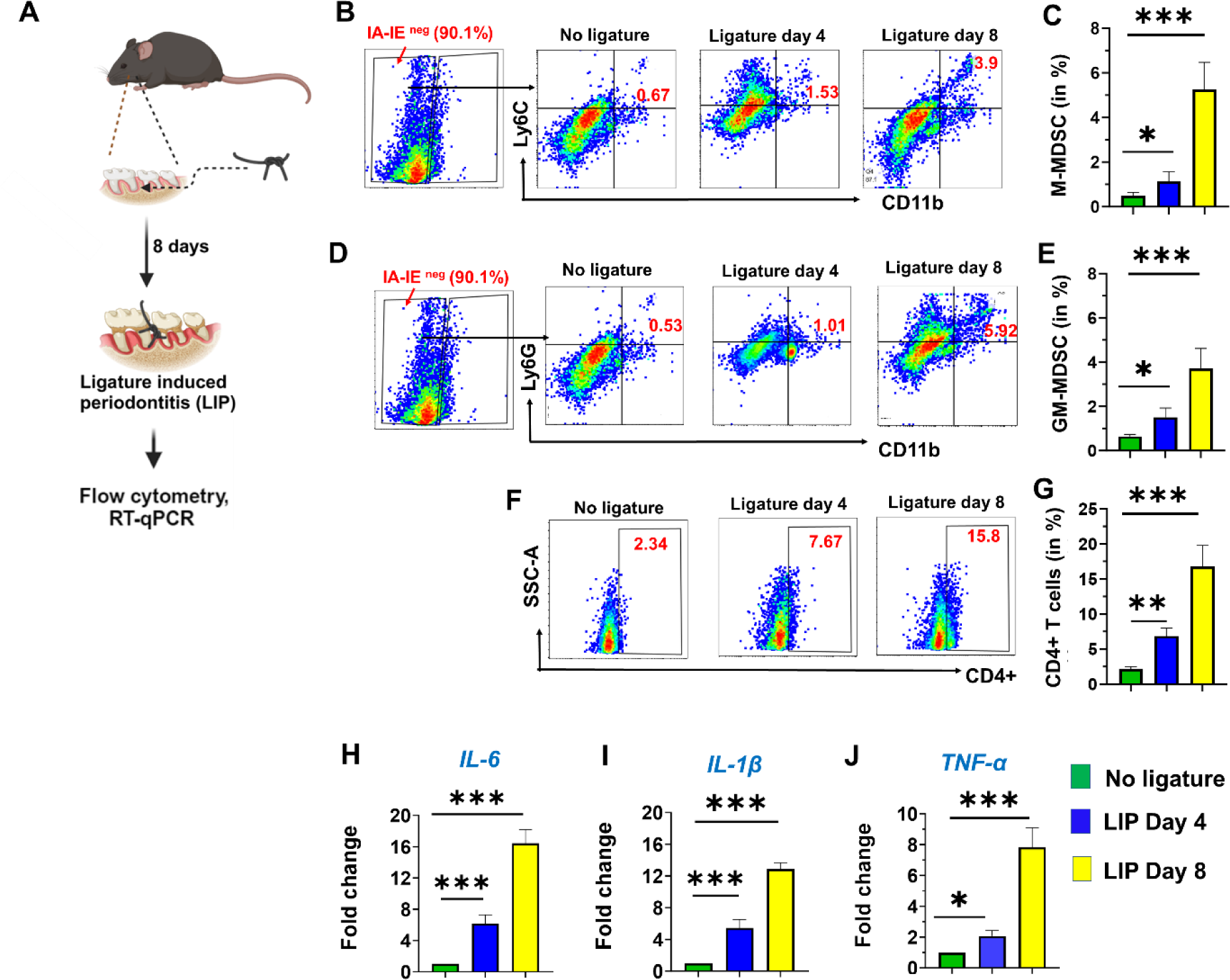
Time-kinetics of MDSC infiltration in a murine model of ligature induced periodontitis (LIP). Mice were subjected to LIP and gingiva were harvested at different time points of disease progression. (A) Schematics of the experiment. Scatter plots showing infiltration of (B) CD11b^+^Ly6C^+^ M-MDSCs and (D) CD11b^+^Ly6G^+^ G-MDSCs in the murine LIP model. Quantitative analysis of (C) M-MDSCs and (E) at different time points during the disease course with respect to no ligature (control mice). (F) Scatter plot showing infiltration of CD4^+^ T cells in gingiva of LIP model during disease progression. (G) Quantitation of infiltrated CD4^+^ T cells in LIP model compared to no ligature. (H) RT-qPCR showing the expression of pro-inflammatory cytokines IL-6, IL-1β and TNFα) in the gingiva derived from no ligature and LIP mice. The Ct values of three replicates were analyzed to calculate fold change using the 2^−ΔΔCt^ method. β-actin was used as a housekeeping control. FlowJo_v10.9.0 was used to create scattered plots in (B), (D) and (F). Each bar represents mean ±S.D. One-way ANOVA was used to calculate the significance in (B), (E) and (G), while Student’s t-test was performed to evaluate the significance in (H). *P< 0.05; **P< 0.01; ***P< 0.001.

Next, we evaluated the presence of pro- and anti-inflammatory cytokines in LIP murine gingiva by RT-qPCR. Our results showed significantly higher expression of pro-inflammatory cytokines IL-6 [fold change: 6.1±1.09 (4DPL) and 16.4±1.72 (8DPL); P<0.001], IL1β [fold change: 4.55±0.65 (4DPL) and 14.45±0.91(8DPL); P<0.001] and TNF-α [fold change: 5.45±1.06 (4DPL) and 12.85±0.82 (8DPL); P<0.001] compared to mice without any ligature (**Fig. 3H**).

### Adoptive Transfer of MDSCs attenuates disease progression in LIP model

MDSC have been shown to promote tissue repair in multiple immune-mediated diseases.^17-19^ MDSC suppress pro-inflammatory T-cell (CD4/CD8) and B cell functions via production of immune-regulatory molecules (IL-10, ARG1, and iNOS).^37,38^ We hypothesize that an imbalance in MDSC maintenance during inflammation will augment the clinical manifestations of PD and asked whether reduced MDSC infiltration or inefficient *in vivo* functional activity is contributory to PD progression. To address this, we performed adoptive transfer of MDSCs to examine their role in PD pathogenesis in our LIP model. **Fig. 4A** shows the schema of our experimental design. After establishing the M- and G-MDSCs populations in the bone marrow by flow cytometry **(Fig. 4B**), we sorted M-MDSCs and G-MDSCs (**Supplementary Fig. 1**). Both subtypes of MDSCs showed suppression of CD4^+^ T cell proliferation in a dose-dependent manner. CFSE^lo^ cell percentage were considered as the proliferating population. The percentage of CD4^+^ T cell proliferation was significantly reduced in the presence of either M-MDSCs or G-MDSCs in higher ratios (40.66 ± 5.14% at 1:10 ratio and 22.18 ± 2.71% at 1:5 ratio) compared to splenocytes + anti CD3/CD28 54.34 ± 3.90% (positive control). Splenocytes alone served as negative control and showed negligible proliferation (0.13 ± 0.04%) (**Supplementary Fig. 2A, 2B**). This observation strongly points to the immunoregulatory nature of murine MDSC infiltrating in gingiva during LIP progression. Next, we infused CFSE labeled M- and G-MDSCs in separate LIP animals via retro-orbital route to restrain PD. Since MDSCs do not undergo proliferation, CFSE dilution was not a limiting factor for their evaluation in gingiva. After 48 h post-delivery, we detected significant percentages of CFSE^+^ cells in the gingiva of LIP mice either infused with M-MDSC (3.16 ± 0.31%) or PMN-MDSC (2.1 ±0.20) populations, which strongly suggests the homing of adoptively transferred MDSCs in murine LIP gingiva (**Fig. 4 C,D**).

**Figure 4.**
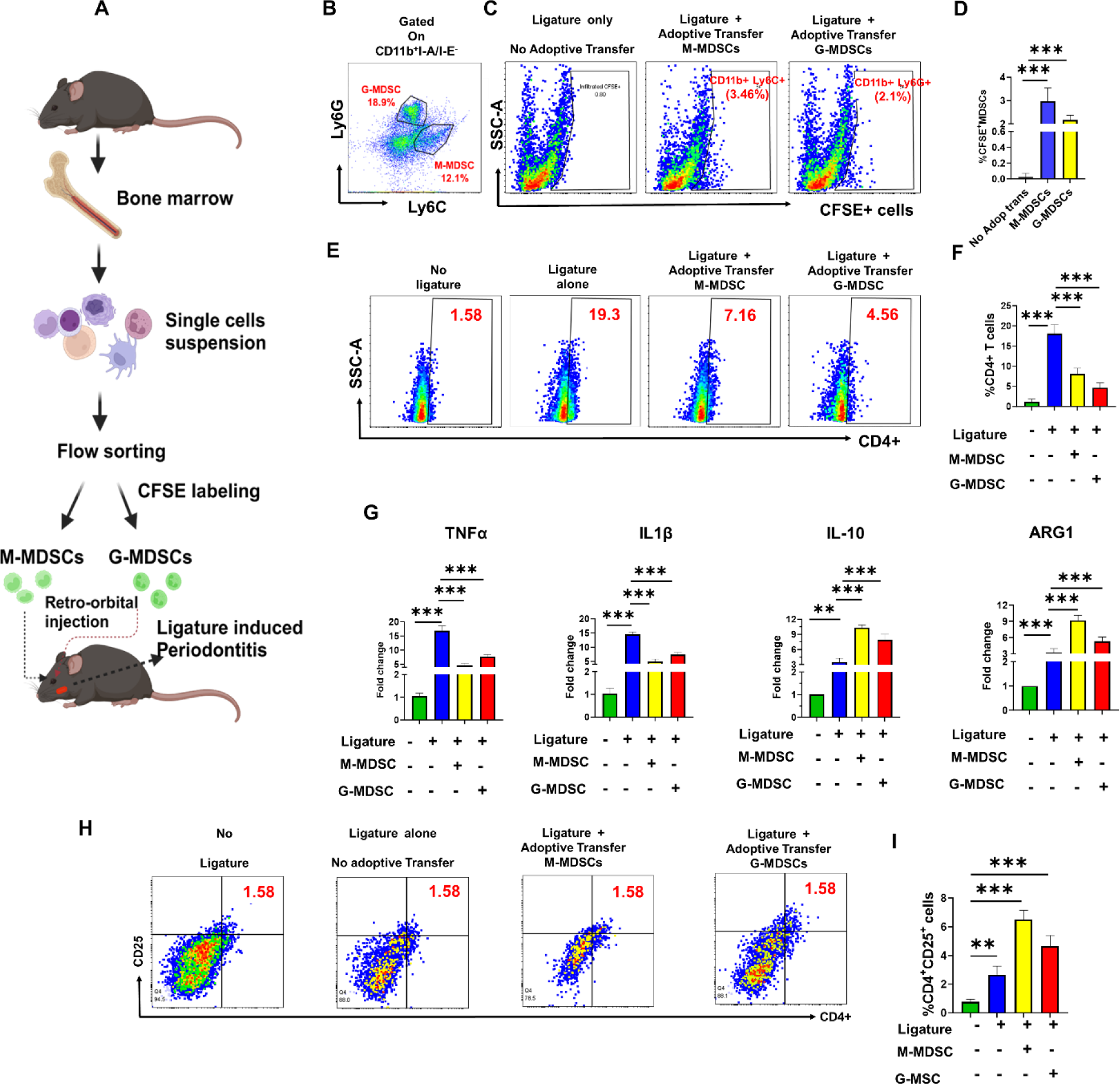
Adoptive transfer of MDSCs inhibits periodontal inflammation in a murine ligature-induced periodontitis model. (A) Overall scheme of the experimental design. (B) Scatter plot showing the staining of M-MDSCs (CD11b^+^I-A/I-E^-^Ly6C^+^) and G-MDSCs (CD11b^+^I-A/I-E^-^Ly6G^+^) cells in mouse bone marrow. CD11b^+^I-A/I-E^-^Gr1+ cells were sorted, CFSE labeled and adoptively transferred (2x10^6^ cells) via retro-orbital route in LIP mice. (C) Scatter plot showing infiltration of bone marrow derived G- and M-MDSCs in mice gingiva 8 days post-delivery. (D) Quantitative analysis of M- and G-MDSCs infiltration in LIP mice gingiva. (C) Percentages of gingiva infiltrated M-MDSC and G-MDSCs in LIP mice. (E) Scatter plot showing the reduced T cell infiltration in mice adoptively transferred with MDSC subsets. (F) Quantitative analysis of gingiva infiltrated CD4^+^ cells in LIP mice undergone adoptive transfer of MDSCs. (G) RT-qPCR of pro-inflammatory (TNFα and IL1β) and anti-inflammatory (IL-10 and ARG1) cytokines in murine gingiva. The Ct values of three replicates were analyzed to calculate fold change using the 2^−ΔΔCt^ method. β-actin was used as a housekeeping control. (G) Flow cytometric analysis of CD4^+^CD25^+^FoxP3^+^ Tregs in LIP model with or without adoptive transfer of MDSCs. (H) Quantitative analysis of Tregs in the gingiva of LIP mice with and without adoptive transfer of MDSCs. N=5 mice for each group was used for the experiment. FlowJo_v10.9.0 was used to create scattered plots in (B), (D) and (G). One-way ANOVA in (C), (E) and (H) and Student’s t-test in (F) were used to calculate P values. *P< 0.05;**P< 0.01; ***P< 0.001.

Next, we adoptively transferred unlabeled 1.5X10^6^ M- or G-MDSCs in mice after ligature placement and monitored their impact on PD progression 8 DPL. Interestingly, significantly reduced CD4^+^ T cell infiltration was observed in LIP mice that underwent adoptive transfer with M-MDSC (8.09±1.42 %; P<0.001) and G-MDSC (4.64±1.17 %; P< 0.001) compared to control (ligature only: 18.1±2.19%) (**Fig. 4 E, F**). Reduced proportion of these cells in gingiva of LIP mice reflect a diminishing degree of periodontal inflammation.^39-42^ To investigate this, we further quantitated the expression of pro-inflammatory (TNF-α and IL-1β) and anti-inflammatory (IL-10) cytokines. Compared to ligature alone, the transcript levels of pro-inflammatory cytokines TNF-α [16.85±1.72% (ligature alone) to 4.5±0.85 % (ligature + M-MDSC) and 7.6±0.83 % (ligature + G-MDSC); P<0.001] and IL-1β [14.5±0.81% (ligature alone) to 4.9±0.90 % (ligature + M-MDSC) and 7.4±0.73 % (ligature + G-MDSC); P<0.001] were significantly reduced at 8 DPL in mice that underwent adoptive transfer of either M-MDSC or G-MDSC. Conversely, expression of anti-inflammatory mediators IL-10 (fold change: 10.35+0.56 (M-MDSC) and 7.9+1.13 % (G-MDSC); P<0.001) and ARG1 (fold change: 9.19+0.95 (M-MDSC) and 5.3+0.79 % (G-MDSC); P<0.001) were significantly increased in mice that received MDSCs (**Fig. 3G**).

Multiple reports have unraveled the ability of MDSCs to facilitate the expansion and recruitment of T regs.^42-45^ Therefore, we examined whether adoptive transfer of MDSCs can increase levels of gingival Tregs. Indeed, we observed significantly higher percentages of CD4^+^CD25^+^ Treg in LIP gingiva + adoptively transferred MDSC [6.4 ± 0.60 % (M-MDSC); P<0.001 and 4.6 ± 0.74 % (G-MDSC); P<0.001] on 8 DPL compared to LIP alone (2.65 ± 0.60 %) and mice without any ligature (control; 0.76 ± 0.18 %). Taken together, these findings clearly suggests that *in vivo* MDSC delivery can robustly mitigate gingival inflammation associated with PD by conferring immunoregulatory function via induction of CD4^+^CD25^+^ cells (**Fig. 4H, I**).

### MDSC depletion accelerates disease severity in a LIP model

To authenticate the above findings, we investigated if gingival depletion of MDSCs will enhance PD progression. Herein, LIP mice were adoptively transferred with I-A/I-E^-^ CD11b^+^Gr1^+^ cells, then treated with the well-known MDSC depletion antibody, anti-Gr1 (clone RB6-8C5)^46,47^ at various time points (0, 2, 4, and 6 DPL) in the buccal and palatal gingiva surrounding the ligature and evaluated the infiltration of MDSCs at 8 DPL. **Fig. 5A** depicts the schematics of the experiment. Compared to ligature alone, we observed significant decreases in M-MDSCs [11.0 ± 1.04 % (0.25 mg/mL), 5.97 ± 1.21 % (0.5 mg/mL) *vs* 16.9 ± 1.42 (Mock)] and G-MDSC populations [2.0 ± 0.31% (0.25 mg/mL) and 0.3 ± 0.29 % (0.5 mg/mL); *vs* 4.7 ± 1.21% (Mock)] in a dose-dependent manner (**Fig. 5B-E)**. Based on the percentage infiltration of MDSCs in gingiva, the effects of anti-Gr-1 were more pronounced for G-MDSCs as compared to M-MDSCs and corroborates with known literature.^48,49^ Next, we examined CD4^+^ T cell infiltration in gingiva and observed a marked dose-dependent increase of CD4^+^T cells in mice treated with anti-Gr-1: 8.9 ± 1.07 % (0.25 mg/mL); 11.1 ± 1.03 % (0.5 mg/mL) and 6.9 ± 0.92 % (Mock). (**Fig. 5F, G**). This data shows that MDSCs tends to embargo CD4^+^ T cell infiltration in gingiva, which has been demonstrated as a necessary event during the progression of periodontitis.^39-42^ Furthermore, compared to ligature, gingival tissues of LIP mice displayed a progressive increase in transcript levels of pro-inflammatory cytokines TNFα [fold change : 13.4 ± 1.83 (Lig) *vs* 17.2 ± 0.56 (0.25 mg/mL and 22.6 ± 1.26 (0.5 mg/mL, P<0.001] and IL1β [fold change: 8.9 ± 1.42 (Lig) *vs* 12.0 ± 1.20 (0.25 mg/mL and 16.3 ± 1.84 (0.5 mg/mL), P<0.001], and concomitant decrease in immunoregulatory genes IL-10 [4.6 ± 0.78 (Lig) *vs* 3.6 ± 0.72 (0.25 mg/mL) and 2.3 ± 0.42 (0.5 mg/mL); P<0.001] and ARG1 [1.93 ± 0.17 (Lig) *vs* 1.34 ± 0.24 (0.25 mg/mL) and 0.62 ± 0.21 (0.5 mg/mL); P<0.001] in a dose-dependent manner (**Fig. 5H**).

**Figure 5.**
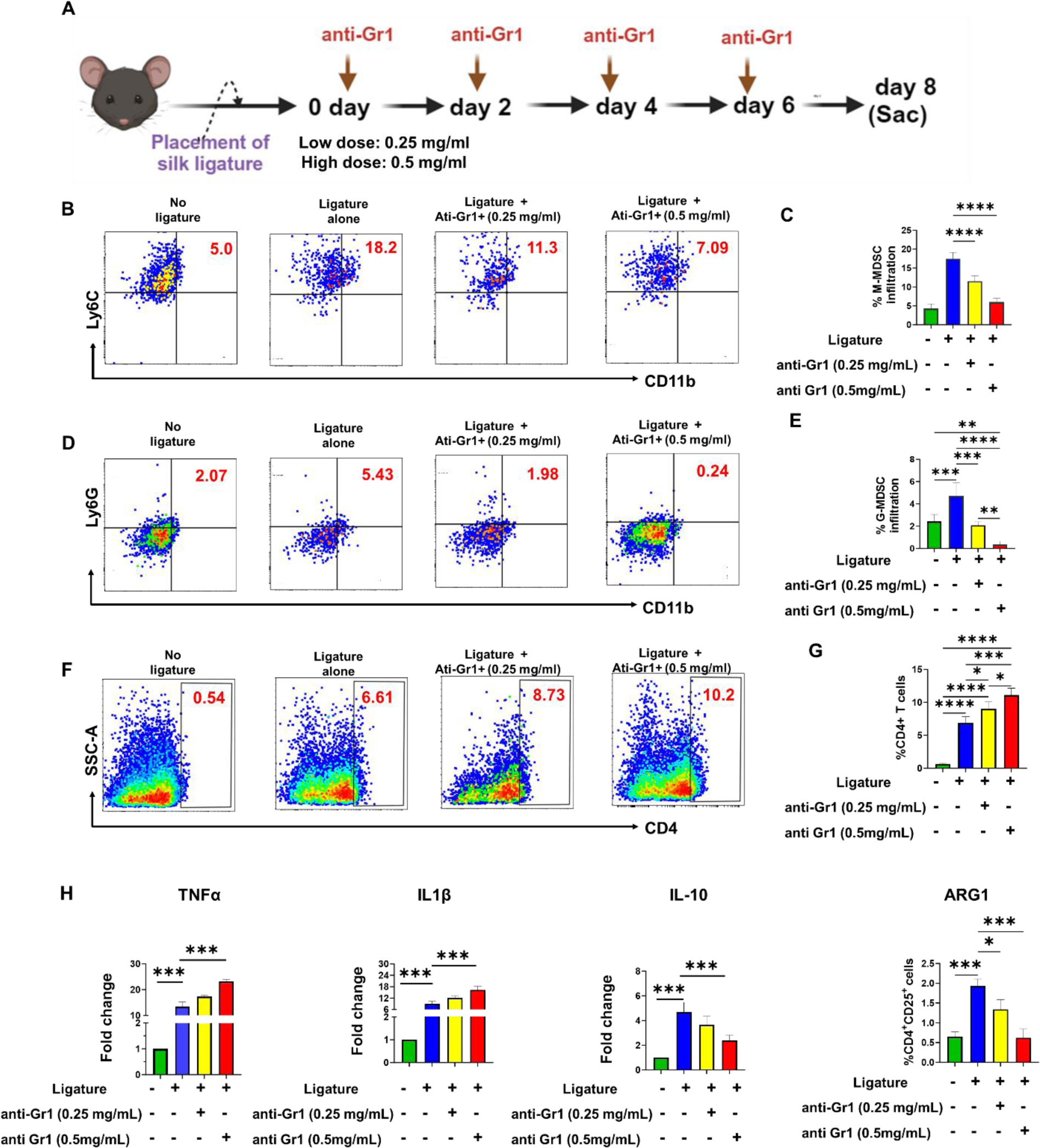
Depletion of MDSCs augments periodontal inflammation. (A) Overall scheme of the experimental design. Mice (n=5/group) were treated with low and high doses of anti-Gr1 antibody or IgG control on alternate days. At day 8, inflammatory status of the gingiva was assessed by various markers by flow cytometry and RT-qPCR. (B) Flow cytometric analysis showing the effect of anti-Gr1 on the infiltration of (B) M-MDSCs and (D) G-MDSCs in the gingiva of murine LIP model. (C) Quantitative analysis of (C) M-MDSCs (CD11b^+^Ly6C^+^) and (E) G-MDSCs (CD11b^+^Ly6G^+^) infiltration at 8 DPL after injecting different doses of anti-Gr1. Higher infiltration of CD4+ T cells occur in MDSC depletion. (F) Scatter plots and (G) quantitative analysis shows significantly higher infiltration of CD4^+^ T cells in the gingiva of LIP mice depleted of MDSCs. (H) RT-qPCR showing the expression of pro-inflammatory (TNFα, IL1β) and anti-inflammatory markers (IL-10, ARG1) in gingiva of LIP mice with/without adoptive transfer of MDSCs. The Ct values of replicates were analyzed to calculate fold change using the 2^−ΔΔCt^ method. β-actin was used as a housekeeping control. Each bar in (C), (E), (G), (F), (H) and (I) represent Mean ±S.D. One-way ANOVA in (C), (E) and (G), while Student’s t-test in (H) were used to calculate P values. *P< 0.05;**P< 0.01;***P< 0.001.

### Adoptive transfer of MDSCs results in attenuated alveolar bone loss

Uncontrolled periodontal inflammation can induce osteoclastogenesis leading to alveolar bone loss. ^50,51^ Therefore, we performed micro-computed tomography (microCT) analysis to evaluate alveolar bone loss in our LIP model. Our results clearly showed progressive alveolar bone loss from 4-to 8 DPL which reflects the presence of an inflammatory microenvironment. This was particularly evident at 8 DPL. We asked whether depletion of MDSC promotes or inhibits osteoclastogenesis. Mice were treated with anti-Gr1 (clone RB6-8C5) antibody to suppress MDSC function and alveolar bone loss quantified by examining the distance from the cementoenamel junction (CEJ) of teeth to the alveolar bone crest (ABC) and % bone volume/total volume (BV/TV), a quantitative indicator of bone mineral density. We measured these parameters in different murine molar teeth as shown in **Fig. 6A** and noted that after 8 DPL, significant bone loss occurred in our LIP mouse model. The CEJ to ABC distance (in µm) in M1A, M1B, and M2 teeth were 20.13 ± 2.4; 14.33 ± 3.06 and 12.10 ± 2.95, respectively. Interestingly, these values increased significantly upon treatment with MDSC depleting antibody (anti-Gr1) (**Fig. 6B**). CEJ to ABC distance in M1A, M2B, and M2 molar teeth were 24.87 ± 1.8, 17.3 ± 2.11. and 14.83 ± 1.22 (in µm), respectively in mice treated with anti-Gr1. Further, a remarkable decrease in % BV/TV was observed under M1 and M2 molars in mice functionally depleted for MDSC compared to ligature only group (**Fig. 6C**). We also noticed significantly lower trabeculae number (3±0.7) in MDSC depleted mice compared to ligature (15±3.5) or no ligature (23±4.2) (**Fig. 6D**) further supporting overall bone loss in the absence of MDSC activity.

**Fig. 6.**
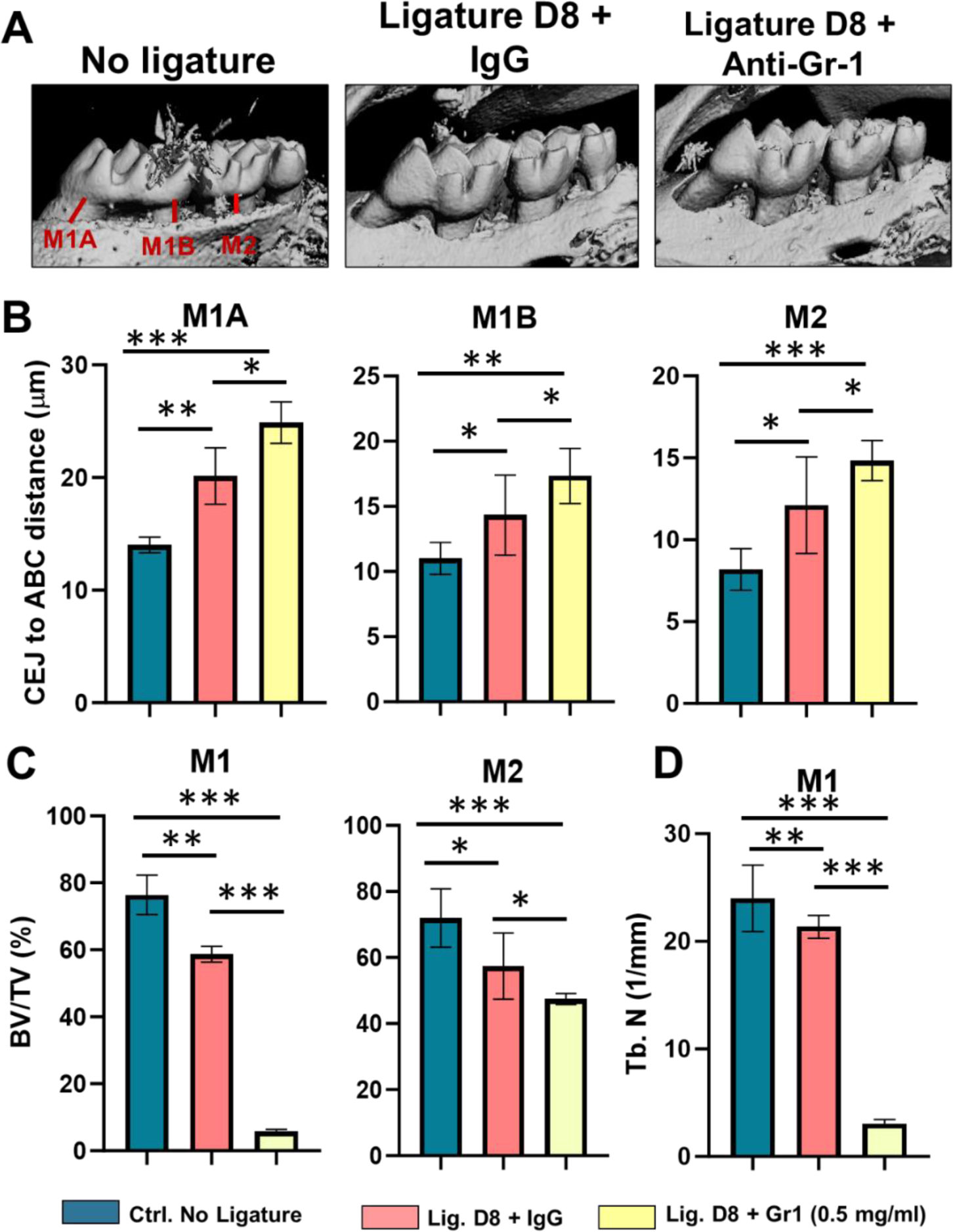
MDSC depletion augments alveolar bone loss in LIP model. Mice were subjected to LIP and treated with IgG, or Anti-Gr1 (0.5 mg/ml) every other day for 8 days by injecting drugs in gingiva using a micro syringe. Animals without ligature were used as controls. Animals were sacrificed at day 8 to examine disease progression by monitoring alveolar bone loss. (A) Representative 3-D microCT images of maxillae after treatments at day 8. (B) Quantitative assessment of alveolar bone loss in mice. The distance (in μm) from the cementoenamel junction (CEJ) to the alveolar bone crest (ABC) at the palatal side was measured at three predetermined maxillary buccal sites shown in the upper panel (M1A, M1B, and M2) by Image J. (C) Histograms showing the bone volume density between root and M1 and between root and M2 at day 8 post-treatment. (D) Trabeculae numbers (/mm) in mice subjected to LIP and treated with or without anti-Gr1. Each bar in (C), (E), (G), (F), (H) and (I) represent Mean ±S.D. One-way ANOVA in (C), (E) and (G), while Student’s t-test in (H) were used to calculate P values. *P< 0.05;**P< 0.01;***P< 0.001

Overall, these observations strongly suggest that depletion of MDSCs, along with their immunomodulatory functions, results in the exacerbation of microbially-induced inflammation and concomitant loss of alveolar bone volume, a hallmark of PD.

## Discussion

Potent immunoregulatory properties of MDSC make them a promising and an attractive therapeutic target for PD treatment; however, their role in PD onset and disease resolution remains elusive. Various reports demonstrate that cell therapy of MDSC, initial responders to tissue damage and facilitators of wound healing, can modulate inflammation and restoring immune homeostasis.^52,53^ The role of MDSCs in periodontal pathogenesis is an understudied topic. In a murine model of obesity and periodontitis, Kwack et al. reported the increase in MDSCs in the spleen and bone marrow of 16 week old mice fed with either a low-fat diet (LFD) or high-fat diet (HFD).^54^ Later, they reported that M-MDSCs derived from HFD fed mice showed an *in-vitro* tendency to form larger osteoclasts.^54^ In agreement, other studies also suggested this narrative noting that in various pathological conditions involving bone destruction, MDSCs can act as osteoclast progenitors.^55-57^ However, to date, no mechanistic data has supported this evidence. Interestingly, Sawant et al., evaluated whether MDSCs from various organs (lung, spleen, blood, lymph nodes) of tumor-challenged mice can differentiate into osteoclast.^55^ Results indicated that none of these MDSCs showed potential to differentiate into osteoclasts, except MDSCs derived from bone of breast cancer mice exhibiting bone metastasis.^55^ This data supports the notion that MDSCs intrinsic potential to convert into osteoclasts is microenvironment specific.

Before exploring MDSCs as a therapeutic tool to ameliorate PD, we asked whether MDSCs home to inflamed gingiva in humans and mice. Here we compared single cell suspension obtained from gingiva derived from healthy and periodontitis subjects. Significantly, a high proportion of HLA-DR^-^CD11b^+^CD33^+^ cells was associated with infiltration of MDSCs during periodontal inflammation. This may be reasoned as an attempt to establish immune tolerance to avoid periodontal tissue damage from excessive gingival inflammation. MDSCs inhibit effector T cells functions via depleting L-arginine through arginase-1 (ARG1), and producing NO, IL-10 and TGF-β.^30.31^ Remarkably high expression of AGR1 and IL-10 in periodontally inflamed gingiva tissues also substantiate MDSC infiltration. Based on the proposed gating strategy, we sorted MDSCs which indeed exhibited *in vitro* suppression of CD4^+^ T cells in a dose dependent manner, thereby substantiating the immune suppressive nature of gingiva infiltrated MDSCs. Furthermore, the above immune suppressive factors pertaining to MDSCs facilitate the recruitment and *de novo* expansion or recruitment of Treg cells.^43-45^ In parallel to MDSCs, Tregs also seem to be increased in gingiva of PD subjects but are unable to resolve the inflammatory environment. We propounded here that numbers of MDSCs in inflamed gingiva were not sufficient to induce needed quantities of Tregs to overwhelm overt inflammation associated with periodontal pathogenesis. Therefore, MDSC therapy may promote resolution of PD. This is the first report clearly demonstrating the infiltration of M- and G-MDSCs in periodontitis subjects.

Next, our ligature induced periodontitis mice model was used to investigate the kinetics of MDSCs infiltration during the progression of PD. Significantly increased infiltration of M-MDSC (HLA-DR^-^CD11b^+^Ly6C^+^) and G-MDSC (HLA-DR^-^CD11b+Ly6G+) cells from day 0-8 DPL further substantiated the progressive increase of MDSC infiltration with the advancement of periodontal inflammation. Furthermore, previous studies on LIP models (either in mouse or rat) were associated with augmented infiltration of CD4+ T cells as well as the increased levels of pro-inflammatory cytokines, IL-1β and TNF-β.^57,58^ In agreement with other studies, we show a significant amount of alveolar bone loss in our murine LIP by microCT image analysis.^59,60^ To evaluate the immunoregulatory nature of M- and G-MDSCs, we co-cultured these cells with splenocytes of the same mice. Similar to human MDSCs, M- and G-MDSCs from gingiva of LIP mice significantly inhibited CD4^+^ T cells proliferation in dose dependent manner.

Collectively, our data authenticates the infiltration of MDSCs with immunoregulatory functions during periodontal pathogenesis. However, questions remain as to why these infiltrating MDSCs were unable to attenuate the progression of periodontal inflammation. In our experiments on T cell proliferation, we have learned that specific doses of MDSCs (G or M) significantly reduce the proliferation of CD4^+^ T cells. Of note, in type 1 diabetic patients, Grohová et al. showed M-MDSC can only suppress the proliferation of autologous T cells in the presence of high MDSC: T cells ratio.^61^ In another study, Fu et al., demonstrated a negative correlation between numbers of MDSC in pancreatic islets and the progression of diabetes. They postulated that reduced levels of islet derived MDSCs was contributory to islet inflammation and failure.^62^ Therefore, we hypothesized that naturally infiltrated MDSCs during disease progression were not in adequate numbers to overcome periodontal inflammation. To test this in our LIP murine model, we performed adoptive transfer of sorted CFSE-labeled I-A/I-E^-^CD11b^+^Gr1^+^ via the retro-orbital route due to its proximity to gingiva. For the first time, our results demonstrated the infiltration of adaptively transferred MDSCs in gingiva via the retro-orbital route. Interestingly, mice gingiva exhibited significantly reduced number of CD4+ T cells (i.e., reduced degree of periodontal inflammation). Furthermore, gingival biopsies of these mice demonstrated comparatively reduced levels of pro-inflammatory cytokines TNFα and IL-1β, and concomitantly increased levels of immune suppressive cytokines, ARG1 and IL-10 post MDSC therapy. Moreover, in I-A/I-E^-^CD11b^+^Gr1^+^ recipient mice, we have shown increased numbers of CD4^+^CD25^+^ Tregs compared to control. This data suggests that adoptive transfer of MDSCs in LIP mice model either induced or facilitated the expansion of Tregs. As indicated by significantly reduced gingival levels of TNFα and IL-1β, this approach may be a therapeutically advantageous, innate pathway aimed at resolution of periodontal inflammation and reversal of disease progression. To further validate the immune protective roles of MDSCs in periodontitis, we injected MDSC-depleting antibody (anti-Gr1) in different doses in our LIP murine model. Interestingly, higher doses of anti-Gr1 significantly reduced the amount of infiltrated MDSCs (both G-MDSCs and M-MDSCs) resulting in increased numbers of CD4^+^ T cells and expression of pro-inflammatory cytokines.

Chronic osteolytic inflammation during periodontitis leads to progressive loss of alveolar bone.^63^ M-MDSC, similar to monocytes, also serve as osteoclast progenitors *in vivo* and may exacerbate disease by enhancing alveolar bone loss; however, it is not clear what percent of infiltrating MDSCs convert to osteoclast precursors.^64-68^ Nonetheless, whether MDSCs directly impact alveolar bone loss has not been investigated in a murine model of PD. Here we demonstrated that adoptive transfer of MDSCs significantly inhibited various pro-inflammatory markers and upregulated expression of anti-inflammatory cytokines suggesting its anti-inflammatory function in periodontal tissues *in vivo*. This is in line with our human *ex vivo* studies showing gingival MDSC isolated from inflamed tissues exhibit immunoregulatory functions as observed by reduced autologous T cell proliferation. Mice depleted of MDSC using anti-Gr1 antibody exhibit higher inflammation further substantiating an immunoregulatory role of MDSC in periodontal tissues and their direct involvement in mitigating inflammation. Importantly, loss of alveolar bone in MDSC-depleted mice as observed by increase in CEJ-ABC distance, lower bone volume and trabeculae numbers indicates severe disease progression in LIP models. These results suggest that MDSCs confer immunoprotection during periodontal pathogenesis and highlight its use as a novel immunotherapeutic target. Overall, our data strongly suggest that immunoregulatory and osteoprotective activity of gingival MDSCs during periodontitis can be leveraged as a potential strategy to ameliorate periodontal inflammation.

## Materials and Methods

### Human Sample collection

This study was approved by the Ethics Committee at The University of Illinois Chicago, College of Dentistry (IRB Protocol# 2017-1064). Subjects presenting to the Postgraduate Periodontics Clinic were recruited for this study. Systemically healthy subjects exhibiting stage III grade B periodontitis or healthy periodontal tissues were recruited for this study as previously described.^29,30^ Periodontitis subjects displayed probing depth ≥ 6mm, clinical attachment loss ≥ 5 mm, bleeding on probing, and radiographic evidence of bone loss. Control healthy periodontal patients displayed probing depths ≤ 3mm, no bleeding on probing, no evidence of clinical attachment loss, and no radiographic evidence of bone loss. Inclusion criteria included male and female subjects ages 18 to 70 years and in good systemic health. Exclusion criteria included chronic disease (diabetes, hepatitis, renal failure, clotting disorders, HIV, etc.), current smokers, antibiotic therapy for any medical or dental condition within a month before screening, and subjects taking medications known to affect periodontal status (e.g. phenytoin, calcium channel blockers, cyclosporine). For periodontally healthy subjects (N=12/group), a single gingival biopsy sample (including gingival epithelium, col area, and underlying connective tissue) was collected at the time of crown-lengthening procedures for esthetic or restorative purposes. For subjects with stage III grade B periodontitis (N=12/group), a similar gingival biopsy sample was collected at the time of periodontal flap surgery. Samples were separately processed for RNA isolation and flow cytometric analysis.

### Murine model of ligature-induced periodontitis

The protocol and procedures were ethically reviewed and approved by the Institutional Animal Care and Use Committee at the University of Illinois Chicago (ACC # 22-122). All experiments were performed in accordance with institutional and national guidelines for the care and use of laboratory animals. Periodontitis was induced using a 6-0 silk ligature placed bilaterally between the maxillary first and second molar around buccal and palatal gingiva) in 8-12-week-old female mice (n=4/group) under anesthesia using intraperitoneal injection of ketamine and xylazine. Animals were sacrificed at 4- and 8-day post-ligature (DPL) placement. Mice with displaced ligature during the experimental period were excluded. Gingival tissues were harvested circumferentially from maxillary first and second molars.

### Gingival cell isolation

Human and murine gingiva were minced with a sterile scalpel and incubated for 1 hour at 37°C for digestion in custom RPMI 1640 containing 1 mg/mL collagenase IV (Millipore Sigma, St. Louis, MO, USA) and 0.1 mg/mL DNase I (Roche Diagnostics, Indianapolis, IN, USA). The cells were filtered using a 70µm cell strainer and 5 mL. DNase I solution (0.1 mg/mL; Millipore Sigma) was added and spun at 800 rpm x 5 min at room temperature. The supernatant was discarded, and cells were incubated twice with DNase I solution at 800 rpm x 5 min and finally resuspended in PBS-1% BSA.

### RNA isolation and RT-qPCR

Total RNA was isolated from gingiva (human and mice) using miRNeasy mini-Kit (Qiagen, Hilden, Germany) according to the manufacturer’s instructions. Approximately 250 ng of total RNA was used to synthesize first-strand of cDNA by Reverse Transcription Kit (Qiagen). For expression analysis of TNFα, IL-1β, IL-10, and ARG1, primers were purchased from Sigma and transcripts were quantified by real-time PCR using a StepOne plus thermocycler (Applied Biosystems, Carlsbad, CA, USA). β-actin was used as housekeeping gene to calculate the fold change expression using delta-delta CT method. The data are presented as normalized fold change: Mean± SD.

### Flow cytometry

Single cell suspension obtained from gingiva (human and murine LIP model) were washed with ice-cold PBS supplemented with 1% (v/v) BSA and 0.08% sodium azide. All samples were stained with the following fluorochrome conjugated antibodies (all from BioLegend, San Diego, CA, USA): anti-human CD11b conjugated with Pacific Blue™ (Clone: ICRF44), anti-human CD33 conjugated with APC (Clone: WM53), anti-human CD14 conjugated with APC-700 (Clone: HCD14), anti-human HLA-DR conjugated with AF488 (Clone: L243), anti-human CD4 conjugated with FITC (Clone: OKT4), anti-human CD25 conjugated with PerCP (Clone: M-A251), anti-human IFN-γ conjugated with APC-Cy7 (Clone: 4S.B3), anti-mouse I-A/IE conjugated with AF488 (Clone: M5/114.15.2), anti-mouse CD11b conjugated with PerCP (Clone: M1/70), anti-mouse Ly6C conjugated with PE (Clone: HK1.4), anti-mouse Ly6G conjugated with APC (Clone: 1A8), anti-mouse CD25 conjugated with APC/Cyanine7 (Clone: 3C7), and anti-mouse Ly-6G/Ly-6C (Gr-1) conjugated with Pacific Blue™ (Clone: RB6-8C5). The antibodies in respective samples were incubated at 4°C for 45 min in PBS-BSA (1%). For intracellular IFN-γ staining, the isolated cells were stimulated with leukocyte activation cocktail in the presence of Golgistop (BD Biosciences, Franklin Lakes, NJ, USA) at 37°C for 4 hours. After surface marker staining, cells were permeabilized using permeabilization buffer (eBioscience, San Diego, CA, USA) and stained with anti-mouse IFN-γ (clone XMG 1.2) antibodies. Cells were then washed two times with PBS supplemented with BSA (1%) and data recorded in a Cytoflex flow cytometer (Beckman Coulter Life Sciences, Indianapolis, IN, USA). Unstained and isotype control-treated cells were used as negative controls. The data was analyzed using FlowJo_v10.9.0 (Ashland, OR, USA).

### Human and murine MDSC sorting and co-culture with respective CD4^+^ T cells

To perform co-culture with human MDSCs, HLA-DR^-^CD11b^+^CD33^+^ both M-MDSCs (HLA-DR^-^CD11b^+^CD33^+^CD14^+^) and G-MDSC (HLA-DR^-^CD11b^+^CD33^+^CD14^-^) were collected by fluorescence-activated cell sorting (FACS). Similarly, MDSCs from gingiva of LIP model were sorted as I-A/I-E^-^ CD11b^+^ Gr1^+^cells. M-MDSC (I-A/I-E^-^ CD11b^+^ Ly6C^+^) and G-MDSCs (I-A/I-E^-^ CD11b^+^Ly6G^+^) comprised the total population of MDSCs. To perform co-culture of human MDSCs, 10^5^ MDSCs were plated in 96 well U-bottom plate and autologous CD4^+^ T cells were added in 1:1, 1:5 and 1:10 ratio keeping the HLA-DR^-^ CD11b^+^CD33^+^ as constant (10^5^ cells) in the presence of ovalbumin peptide (1 mg/mL). CD11b^+^HLA-DR^+^ myeloid cells were added in the same proportion as MDSCs for antigen presentation. To test the immunoregulatory nature of murine gingiva derived MDSCs, I-A/I-E^-^CD11b^+^Gr1^+^cells were plated in 96 well U bottom plates (10^5^) and autologous splenocytes were added in 1:1, and 1:10 ratio. Prior to culture, human CD4^+^ T cells and splenocytes were labeled with CFSE (10μM; ThermoFisher, Waltham, MA, USA), according to manufacturer’s instructions. The co-cultures were harvested at day 5 and cells labeled with anti-anti human or mouse CD4 for 45 min on ice in PBS-1% BSA, washed twice in PBS-1% BSA (w/v) and data acquired in a Cytoflex flow cytometer and analyzed by FlowJo_v10.9.0.

### Adoptive transfer of MDSCs

Spleens were processed by mechanical trituration following red blood cell lysis. Cell suspensions were filtered through a 70 µm strainer, washed twice with PBS. Finally, splenocytes were stained with fluorescence-conjugated anti-mouse antibodies for cell surface markers, including CD11b (clone M1/70), Gr-1 (clone RB6-8C5) and Ly6G (clone: S19018G) or Ly6C (clone: RB6-8C5), and sorted via FACS. Using a micro syringe, 1 x10^6^ flow-sorted I-A/I-E^-^CD11b^+^Gr1^+^ cells were resuspended in saline and infused into the retro-orbital space (behind the eye globe) in the anesthetized mice after ligature placement. Control group animals were injected with saline alone. Animals were humanely euthanized after 8 DPL. We also tested low (0.5 mg/mL) and high doses (1 mg/mL) of MDSC depletion antibody, anti Gr-1, to block the infiltration and activity of MDSCs in the gingiva by local injection.

### microCT analysis

The maxillae were separated and fixed in 4% neutral paraformaldehyde for 24 h and stored in 70% ethanol at 4 °C till further analysis. The key parameters of the micro-CT scanner (Siemens Medical Solutions USA, Inc., Malvern, PA, USA) were as follows: source voltage of 60 kV, current of 220 μA, and resolution of 8.89 μm. Three-dimensional (3D) images were measured by the Inveon Research Workshop software (Siemens Medical Solutions USA, Inc.). For linear measurements, the height of alveolar bone loss was measured as the distance from the cementoenamel junction (CEJ) to the alveolar bone crest (ABC). The buccal groove, palatal groove, mesio-buccal, disto-buccal, mesio-palatal, and disto-palatal sites of the maxillary second molars were chosen as anatomical measurement sites. All data of the CEJ -ABC distance at mentioned sites were analyzed by three blinded investigators. We also qualitatively examined other alveolar bone measures including percent bone volume (bone volume/tissue volume, BV/TV) and trabecular number (Tb.N). For volumetric measurement, the amount of circumferential bone loss of maxillary second molars was assessed following the protocol described previously.^63^ Briefly, contours were drawn beginning below the roof of the furcation and moving in the apical direction until the first appearance of the alveolar bone crest. The most distal root of the first molar and the most mesial root of the third molar were used as borders. All contours were drawn at regular intervals (every five data planes).

### Statistical analysis

All the data were analyzed using GraphPad Prism (GraphPad Software, La Jolla, CA, USA) and presented as Mean ±SD. P values were calculated using either a one-way ANOVA or Student t test.

## AUTHOR CONTRIBUTIONS

Study design/funding: AN; data acquisition: RN and AV, data analyses: RN, AV and AN; writing manuscript: RN, AV, SN, TVD and AN; all authors reviewed and approved of the final version of the manuscript.

## References

1. Dye, B.A. Global periodontal disease epidemiology. Periodontol. 2000. 58, 10–25 (2012).

2. Albandar, J.M. Epidemiology and risk factors of periodontal diseases. Dent. Clin. North Am. 49, 517–32 (2005).

3. American Academy of Periodontology. 2000. Parameter on chronic periodontitis with slight to moderate loss of periodontal support. J Periodontol. 71, 853–855 (2000).

4. Morelli, T. et al. Periodontal profile classes predict periodontal disease progression and tooth loss. J Periodontol. 89, 148–156 (2018).

5. Nares, S. et al. Rapid myeloid cell transcriptional and proteomic responses to periodontopathogenic Porphyromonas gingivalis. Am. J Pathol. 174, 1400–14014 (2009).

6. Seitz, M. W. et al. Current knowledge on correlations between highly prevalent dental conditions and chronic diseases: An umbrella review. Prev. Chronic Dis. 16, E132 (2019).

7. Sparks, S. P. et al. Serum antibodies to periodontal pathogens are a risk factor for Alzheimer’s disease. Alzheimers Dement. 8, 196–203 (2012).

8. de Molon, R. S. et al. Linkage of Periodontitis and rheumatoid arthritis: Current evidence and potential biological interactions. Int. J Mol. Sci. 20, E4541 (2019).

9. Beck, J. D. & Offenbacher, S. Systemic effects of periodontitis: epidemiology of periodontal disease and cardiovascular disease. J Periodontol. 76, 2089–2100 (2005).

10. Cekici, A., Kantarci, A., Hasturk, H. & Van Dyke, TE. Inflammatory and immune pathways in the pathogenesis of periodontal disease. Periodontol 2000. 64, 57–80 (2014).

11. Lam, R.S. et al. Macrophage depletion abates *Porphyromonas gingivalis*-induced alveolar bone resorption in mice. J Immunol. 193, 2349–2362 (2014).

12. Jotwani, R. & Cutler, C.W. Multiple dendritic cell (DC) subpopulations in human gingiva and association of mature DCs with CD4+ T-cells in situ. J Dent Res. 82, 736–741(2003).

13. Hasturk, H. & Kantarci, A. Activation and resolution of periodontal inflammation and its systemic impact. Periodontol. 2000. 69, 255–273 (2015).

14. Hajishengallis, G. Immunomicrobial pathogenesis of periodontitis: keystones, pathobionts, and host response. Trends Immunol. 35, 3–11(2014).

15. Mombelli, A. Microbial colonization of the periodontal pocket and its significance for periodontal therapy. Periodontol 2000. 76, 85–96 (2018).

16. Gabrilovich, D. I. & Nagaraj, S. Myeloid-derived suppressor cells as regulators of the immune system. Nat. Rev. Immunol. 9,162–174 (2009).

17. Gabrilovich D. I. Myeloid-derived suppressor cells. Cancer Immunol. Res. 5, 3–8 (2017).

18. Tesi, R. J. MDSC; the most important cell you have never heard of. Trends Pharmacol Sci. 40, 4–7 (2019).

19. Bronte, V. et al. Recommendations for myeloid-derived suppressor cell nomenclature and characterization standards. Nat. Commun. 7,12150 (2016).

20. Luker, A. J. et al. The DNA methyltransferase inhibitor, guadecitabine, targets tumor-induced myelopoiesis and recovers T cell activity to slow tumor growth in combination with adoptive immunotherapy in a mouse model of breast cancer. BMC Immunol. 21, 8 (2020).

21. Poschke, I. & Kiessling, R. On the armament and appearances of human myeloid-derived suppressor cells. Clin. Immunol. 144, 250–268 (2012).

22. Dumitru, C. A. et al. Neutrophils and granulocytic myeloid-derived suppressor cells: immunophenotyping, cell biology and clinical relevance in human oncology. Cancer Immunol. Immunother. 61,1155–1167 (2012).

23. Mandruzzato, S. et al. Toward harmonized phenotyping of human myeloid-derived suppressor cells by flow cytometry: results from an interim study. Cancer Immunol. Immunother. 65, 161–169 (2016).

24. Kwak, Y., Kim, H. E. & Park, S. G. Insights into myeloid-derived suppressor cells in inflammatory diseases. Arch. Immunol. Ther. Exp. (Warsz). 63, 269–285 (2015).

25. Yang, H. et al. Myeloid-derived suppressor cells in immunity and autoimmunity. Expert Rev. Clin. Immunol. 11, 911–919 (2015).

26. Ma, H. & Xia, C. Q. Phenotypic and functional diversities of myeloid-derived suppressor cells in autoimmune diseases. Mediators Inflamm. 4316584 (2018).

27. Zhang, Q., Fujino, M., Xu, J. & Li, X. K. The role and potential therapeutic application of myeloid-derived suppressor cells in allo- and autoimmunity. Mediators Inflamm. 421927 (2015).

28. Koehn, B. H. & Blazar, B. R. Role of myeloid-derived suppressor cells in allogeneic hematopoietic cell transplantation. J Leukoc. Biol. 102, 335–341 (2017).

29. Barbour, S. E., et al. Tew JG. Monocyte differentiation in localized juvenile periodontitis is skewed toward the dendritic cell phenotype. Infect. Immun. 70, 2780–2786 (2002).

30. Holmgaard, R. B. et al. Tumor-expressed ido recruits and activates MDSCs in a treg-dependent manner. Cell Rep. 13, 412–424 (2015).

31. Munn, D. H. Blocking IDO activity to enhance anti-tumor immunity. Front. Biosci. **E4**, 734–745 (2012).

32. Campbell, L., Millhouse, E., Malcolm, J. & Culshaw S. T cells, teeth and tissue destruction—what do T cells do in periodontal disease? Mol. Oral. Microbiol. 31, 445–456 (2016).

33. Berglundh, T., Liljenberg, B., Tarkowski, A. & Lindhe, J. The presence of local and circulating autoreactive B cells in patients with advanced periodontitis. J. Clin. Periodontol. 29, 281–286 (2002).

34. Uttamani, J. R, et al. Dynamic changes in macrophage polarization during the resolution phase of periodontal disease. bioRxiv. Preprint. 2023 Feb 21.

35. Ahmad, I., Naqvi, R. A., Valverde, A., & Naqvi, A. R. LncRNA MALAT1/microRNA-30b axis regulates macrophage polarization and function. Front Immunol. 4, 1214810 (2023).

36. Peranzoni, E. et al. Myeloid-derived suppressor cell heterogeneity and subset definition. Curr. Opin. Immunol. 22, 238–244 (2010).

37. Fujii, W. et al. Myeloid-derived suppressor cells play crucial roles in the regulation of mouse collagen-induced arthritis. J Immunol.191,1073–1081 (2013).

38. Dorhoi, A. & Du Plessis, N. Monocytic myeloid-derived suppressor cells in chronic infections. Front Immunol. 4, 1895 (2018).

39. Ben-Sasson, S.Z. et al. IL-1 acts on CD4 T cells to enhance their antigen-driven expansion and differentiation. Proc. Natl. Acad. Sci. USA, 106, 7119–7124 (2009).

40. Gonzales, J.R., Groeger, S., Johansson, A., & Meyle, J. T helper cells from aggressive periodontitis patients produce higher levels of interleukin-1 beta and interleukin-6 in interaction with *Porphyromonas gingivalis*. Clin. Oral Investig. 18, 1835–1843 (2014).

41. Taubman, M. A. & Kawai, T. Involvement of T-lymphocytes in periodontal disease and in direct and indirect induction of bone resorption. Crit. Rev. Oral Biol. Med. 12:125–135 (2001).

42. Alvarez, C. et al. RvE1 impacts the gingival inflammatory infiltrate by inhibiting the t cell response in experimental periodontitis. Front Immunol. 3, 12:664756 (2021).

43. Fujimura, T. et al. Regulatory T cells stimulate B7-H1 expression in myeloid-derived suppressor cells in ret melanomas. J Invest. Dermatol. 132, 1239–1246 (2012).

44. Serafini, P., Mgebroff, S., Noonan, K., & Borrello, I. Myeloid-derived suppressor cells promote cross-tolerance in B-cell lymphoma by expanding regulatory T cells. Cancer Res. 68, 5439–5449 (2008).

45. Schlecker, E. et al. Tumor-infiltrating monocytic myeloid-derived suppressor cells mediate CCR5-dependent recruitment of regulatory T cells favoring tumor growth. J Immunol. 189, 5602–5611 (2012).

46. Ma, C. et al. Anti-Gr-1 antibody depletion fails to eliminate hepatic myeloid-derived suppressor cells in tumor-bearing mice. J. Leukoc. Biol. 92, 1199–1206 (2012).

47. Srivastava, M.K. et al. Myeloid suppressor cell depletion augments antitumor activity in lung cancer. PLoS ONE.;7:e40677 (2012).

48. Ma, C. & Greten, T.F. Editorial: “Invisible” MDSC in tumor-bearing individuals after antibody depletion: Fact or fiction? J. Leukoc. Biol. 99, 794 (2016).

49. Xing, Y. F. et al. Issues with anti-Gr1 antibody-mediated myeloid-derived suppressor cell depletion. Ann. Rheum Dis. 75, e49 (2016).

50. Hajishengallis, G. Periodontitis: From microbial immune subversion to systemic inflammation. Nat. Rev. Immunol. 15, 30–44 (2015).

51. Graves, D. T., Li, J. & Cochran, D. L. Inflammation and uncoupling as mechanisms of periodontal bone loss. J Dent. Res. 90, 143–53 (2011).

52. Mahdipour, E., Charnock, J. C. & Mace KA, Hoxa3 promotes the differentiation of hematopoietic progenitor cells into proangiogenic Gr-1+CD11b+ myeloid cells, Blood. 117, 815–826 (2011).

53. Ou, L. et al. Kruppel-like factor KLF4 facilitates cutaneous wound healing by promoting fibrocyte generation from myeloid-derived suppressor cells. J Invest. Dermatol. 135, 1425–1434 (2015).

54. Kwack, K. H. et al. Novel preosteoclast populations in obesity-associated periodontal disease. J Dent. Res. 101, 348–356 (2022).

55. Sawant, A., & Ponnazhagan, S. Myeloid-derived suppressor cells as osteoclast progenitors: a novel target for controlling osteolytic bone metastasis. Cancer Res. 73, 4606–4610 (2013).

56. Su, L. et al. Phenotype and function of myeloid-derived suppressor cells induced by Porphyromonas gingivalis infection. Infect. Immun. 85, e00213–17 (2017).

57. Zhang, H. et al. Myeloid-derived suppressor cells contribute to bone erosion in collagen-induced arthritis by differentiating to osteoclasts. J Autoimmun. 65, 82–89 (2015).

58. Lin, P. et al. Application of ligature-induced periodontitis in mice to explore the molecular mechanism of periodontal disease. Int. J Mol. Sci. 18, 8900 (2021).

59. Li, C.H., & Amar, S. Morphometric, histomorphometric, and microcomputed tomographic analysis of periodontal inflammatory lesions in a murine model. J. Periodontol. 78,1120–1128 (2007).

60. Yoon, H. et al. Temporal changes of periodontal tissue pathology in a periodontitis animal model. J Periodontal Implant Sci. 53, 248–258 (2023).

61. Grohová, A. et al. Myeloid - derived suppressor cells in Type 1 diabetes are an expanded population exhibiting diverse T-cell suppressor mechanisms. PLoS One. 15, e0242092 (2020).

62. Fu, W. et al. Early window of diabetes determinism in nod mice, dependent on the complement receptor crig, identified by noninvasive imaging. Nat. Immunol. 13, 361–368 (2012).

63. Yu, B., Wang, C.Y. Osteoporosis and periodontal diseases – An update on their association and mechanistic links. Periodontol. 2000. 1: 99–113.(2022).

64. Udagawa, N., Takahashi, N., Akatsu, T. et al. Origin of osteoclasts: mature monocytes and macrophages are capable of differentiating into osteoclasts under a suitable microenvironment prepared by bone marrow-derived stromal cells. Proc. Natl. Acad. Sci. U S A. 87: 7260-4 (1990).

65. Gabrilovich, D. I. Myeloid-Derived Suppressor Cells. Cancer Immunol. Res. 5: 3–8 (2017).

66. Tesi, R. J. MDSC; the Most Important Cell You Have Never Heard Of. Trends Pharmacol. Sci. 40: 4–7 (2019).

67. Bronte, V., Brandau, S., Chen, S. H. et al. Recommendations for myeloid-derived suppressor cell nomenclature and characterization standards. Nat. Commun. 7:12150 (2016).

68. Fujii W, Ashihara E, Hirai H. et al. Myeloid-derived suppressor cells play crucial roles in the regulation of mouse collagen-induced arthritis. J Immunol.;191(3):1073–81 (2013).

