## Supplementary Figures for "Myeloid-derived Suppressor Cells Mitigate Inflammation in Periodontal Disease"

*Supplemental Data*

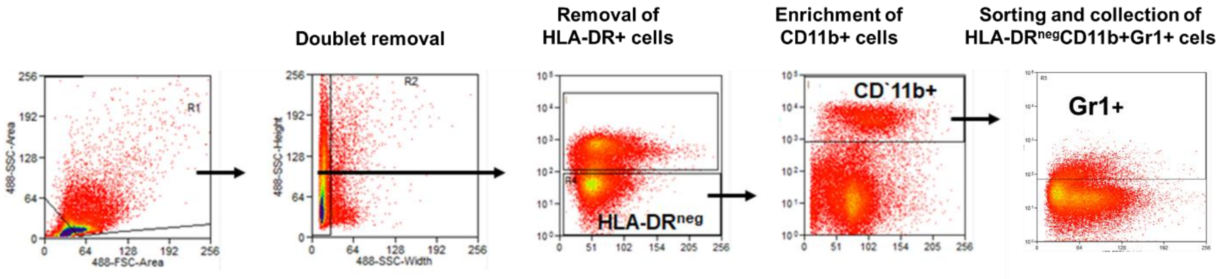


***Supplementary Fig. 1**. **Flow sorting of granulocytic (G)- and monocytic (M)-MDSCs from mice spleen**. After the processing of spleen, RBC were lysed using ACK lysis buffer and the cells were filtered (70-mm strainer) and washed two times with 1x PBS. Splenocytes were stained with CD11b, HLA-DR, and Gr1 antibodies at 4°C for 30 min. Cells were washed with PBS-1% BSA followed by flow sorting. HLA-DR- cells were interrogated for the expression of CD11b and Gr1 (which contains both Ly6G and Ly6C) for sorting. The sorted cells were collected in complete RPMI (10% FBS, 1% Pen-Strep).*

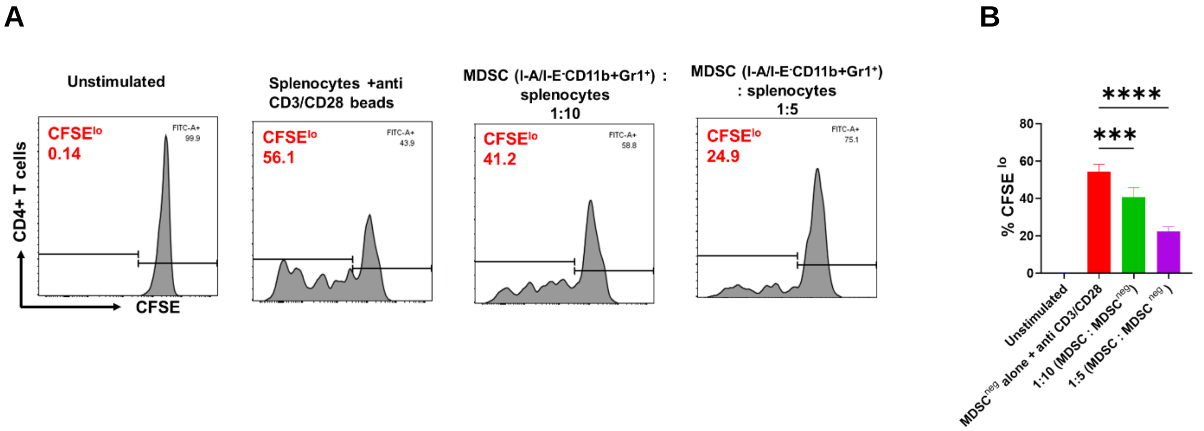


***Supplementary Fig 2**: **Suppression of CD4+ T cells proliferation via sorted murine MDSCs (I-A/I-E$^-$CD11b+Gr1$^+$) in a dose-dependent manner.*** *Splenocytes were stained with CFSE prior to coculture as described in material and methods and then cultured with MDSC in 1:5 and 1:10 ratio. As a positive control proliferation, we used mouse specific anti CD3/CD28 beads (Miltenyi). After five days the co-culture were kept at 4º C and the cells were stained with anti-mouse CD4 (Biolegend) for 45 min at ice and then washed three times with PBS-1%BSA. CFSE$^{hi}$ cells show non-proliferation fraction while CFSE$^{lo}$ exhibit actively proliferating cells. These cells were first gated on CD4+ cells and then CFSE$^{hi}$ or CFSE$^{lo}$ fraction was evaluated. (A) Scatter plots showing the proliferation of CD4+ T cells in mouse splenocytes. (B) Overall CD4+ proliferation reduction in MDSC treated cocultures. This data was analyzed in FlowJO_v_10.9.2. One-way ANOVA was used to calculate the significance. *P< 0.05;**P< 0.05;***P< 0.005;****P< 0.0005.*
